# Female Bengalese finches recognize their father’s song as sexually attractive

**DOI:** 10.1101/2021.07.23.453535

**Authors:** Tomoko G. Fujii, Kazuo Okanoya

## Abstract

Birdsong is an important communication signal used in mate choice. In some songbirds, only males produce songs while females do not. Female birds are sensitive to inter- and intra-species song variation. Some aspects of female song preference depend on developmental experiences. For example, in Bengalese finches and zebra finches, adult females prefer the song to which they were exposed early in life, such as the father’s song. However, it is unclear whether such song preference in females is sexually motivated. The purpose of our study is to test if female Bengalese finches recognize their father’s song as sexually attractive. We measured copulation solicitation displays during playbacks of the father’s song vs. unfamiliar conspecific songs and found that across individuals, the father’s song elicited more displays than other songs. In addition, we analyzed if a bird’s response to a given song could be predicted by the level of similarity of that song to the father’s song. The results suggest that preference for the father’s song in this species is actually relevant to mate choice. Although more precise control is necessary in future studies to elucidate the process of preference development, our results imply the significance of early-life experience in shaping female song preference.

## Introduction

Song is an important communication signal used for mate choice in songbirds [1]. In some species of songbirds, only males produce songs. While females do not sing, they can perceive inter- and intra-species song variations and change their behavior depending on features of the song that reflect the sex, species, and condition of the signaler [2]. For example, female birds selectively respond to conspecific songs over heterospecific ones [3–5]. In some species, this preference seems to be a common tendency across individuals that develops independent of experience after hatching [6,7]. However, developmental experience plays a significant role in shaping other aspects of female song preference. A cross-fostering study in the zebra finch (*Taeniopygia guttata*) demonstrated that recognition of a subspecies based on song depends on auditory experience with the foster father’s song [8]. Early-life song exposure in this species is also critical to be able to discriminate song quality [7] or song performance related to social context [9].

In addition, developmental experience might be an important source of individual differences in song preference. Previous studies in zebra finches and Bengalese finches (*Lonchura striata* var. *domestica*) found that females prefer songs to which they were exposed early in life, such as the father’s song [10–15]. These studies clearly show that female birds acquire and retain an ability to discriminate their (foster) father’s song from other songs, but it is not clear whether such song preference is sexually motivated. Although the possibility of sexual imprinting on the father’s song has already been discussed in some literature [10,14], the idea has not yet been directly tested. Because in these previous studies preference was measured by behaviors such as approach to sound sources [10,11,15] or operant behaviors associated with song playback [13,14], the results could be interpreted as either general selectivity to a familiar stimulus or as a sexual response in the context of mate choice. Likewise, from the perspective of adaptation, it is not clear if preference for a trait similar to that of the parents actually increases an animal’s fitness [16–18]. Some studies in songbirds and other bird taxa indicate that memorizing the traits of parents is a possible strategy for kin recognition to avoid inbreeding [19,20]. Therefore, elucidating why birds prefer their father’s song over unfamiliar conspecific songs is important for a deeper understanding of female mate choice.

In the current study, we aimed to test whether adult female Bengalese finches recognize their father’s song as a sexually attractive signal. To meet this purpose, we measured copulation solicitation displays (CSDs) in response to song playback. CSD is a typical posture that females perform when they accept male courtship, and researchers have utilized it as a reliable index of sexual motivation in response to songs [21–23]. We also analyzed if song preference in our subjects could be predicted by the similarity of the stimulus to the father’s song.

## Methods

### Animals

We used 10 adult female Bengalese finches as subjects for the song preference test. They were obtained from 8 different clutches in our laboratory colony. For breeding, 15 adult finches were used (7 males and 8 females; one male was paired with 2 different females successively). The 15 adults were either bred in our laboratory or purchased from a commercial breeder. The female subjects were raised by both parents and housed with their families (parents and siblings) in a home cage (size: 30 cm wide × 24 cm deep × 33 cm high) until approximately 120 days post hatch (dph). Each home cage was placed in a colony room but visually isolated from one another. The female subjects were kept in a single-sex group cage (37 × 42 × 44 cm) after separation from the parents and male siblings (if any). The number of birds kept in a group cage ranged from 6 to 14. All subjects were sexually mature at the beginning of the experiment (mean age = 279 dph, range = 164-376 dph). No birds had experienced breeding prior to the current study. For the preparation of song stimuli, 19 adult (> 180 dph) male Bengalese finches were used, 7 of which were fathers of the subjects described above.

All birds were kept under a 14:10 h light:dark cycle with food and water provided ad libitum. They were also fed oyster shell and greens once a week. The temperature and humidity were maintained at around 25 oC and 60%, respectively. After finishing all the experiments, the subjects, their parents, and the birds used for stimulus recordings continued to be housed in our colony for additional research. All experimental procedures in this study were approved by the Institutional Animal Care and Use Committee at the University of Tokyo (permission #27-9 and #2020-2).

### Song preference test

Briefly, we designed a playback experiment to test if females perform more CSDs to their father’s song compared to unfamiliar conspecific songs. The overall schedule of the experiment is as follows: the birds first received a hormone implant, then they were moved to the experimental environment for acclimation and isolation from males, and finally they were tested with song playback (Fig. 1(a)).

**Fig 1.**
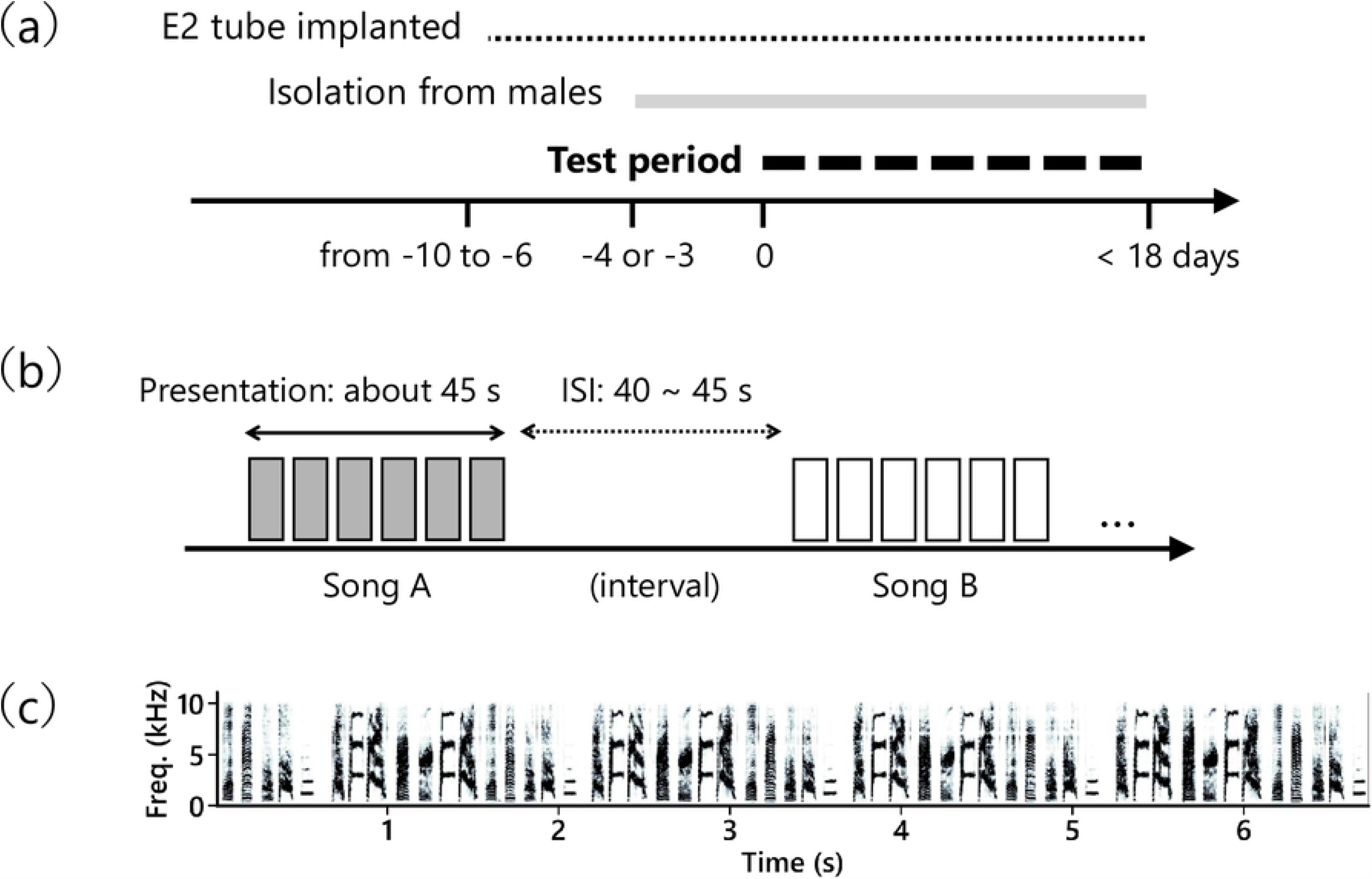
Experimental schedule and stimuli. **(a)** Overall schedule of the experiment. Each line above the horizontal axis indicates the period in which birds were under that manipulation. The start of the song playback test was set as day zero. **(b)** A schematic diagram of the stimulus presentation schedule in a single test. Five songs, each composed of 6 renditions (6 rectangles in the figure) were played. **(c)** A spectrogram of one rendition of the song stimulus is shown as an example.

### Hormone implantation surgery

Female songbirds seldom perform CSDs in response to song playback alone in the laboratory, but studies have shown that subcutaneous implantation of estradiol increases this behavior [21]. Thus, we adopted this hormone implantation method in our experiment and confirmed that the method did indeed increase CSDs in response to song playback (S1 File). Following a previous study using the same bird species [22], we administered 17β-estradiol (E2758, Sigma Aldrich) in silastic tubing with a 1.0 mm inner diameter (100-0N, Kaneka medical products) containing 8 mm of hormone in powder form. During the surgery, a bird was manually restrained, and lidocaine (Xylocaine, AstraZeneca) was applied onto the skin for local anaesthesia. A small incision was made in the skin on the bird’s back and a tube filled with hormone was inserted subcutaneously. The test period started 6 – 10 days after the surgery (Fig. 1(a)). After finishing all the tests, local anaesthesia was again applied to the bird’s back and the hormone tube was excised. From the day of implantation, we continued to monitor the physical condition of the birds every day until a few days after the tube removal.

### Preparation and presentation of song stimuli

In the song preference tests, we presented 5 different conspecific songs to each subject: the father’s song and 4 unfamiliar songs. Females had never been exposed to these 4 unfamiliar songs prior to the experiment. We presented the song of one subject’s father to 2 or 3 other subjects (that hatched in a different clutch) to exclude the possibility that a particular characteristic of a given song generally attracts females, independent of experience.

To prepare stimuli, we recorded songs from adult male Bengalese finches in a soundproof chamber through a microphone (PRO35, Audio-technica) fixed at the top of the cage. The signal from the microphone was amplified (QuadMic, RME) and digitized (Delta66, M-AUDIO) at 16 bits with a 44.1 kHz sampling rate. All recordings were undirected songs, as each bird was kept alone in the chamber during recording. For each male singer, we randomly selected 6 renditions of songs that were recorded without any background noise (Fig. 1(c)). The duration of a single song rendition was 7.04 ± 0.64 (mean ± *s*.*d*.) seconds. The sound waveform was first band-pass filtered at 0.5 – 10 kHz. We normalized the sound amplitude by the standard deviation of the waveform. For each singer, a song was composed of 6 song renditions presented with a 700-ms interval between each rendition (thus approximately 45 seconds in total, Fig. 1(b)). The order of the 6 renditions within a song was randomized each time. Song stimuli were played back from a loudspeaker (MM-SPL2N2, SANWA Supply) positioned next to the test cage. The equivalent continuous A-weighted sound pressure level was measured at a point in the test cage that was 22.5 cm away from the speaker and adjusted to approximately 67 dB using a sound level meter (NL-27, Rion).

### Environment and procedure for song preference tests

Preference tests were conducted in a test cage (30 × 24 × 33 cm) placed in a sound attenuation room (163 × 163 × 215 cm). The light:dark cycle, temperature, and humidity of the testing environment was the same as in the colony room. Subjects were moved to the test cage in a group of either 3 or 4 birds (10 birds were divided into 3 groups) at least 3 days prior to the first test. The purpose of this was to acclimate the birds to the testing environment as well as to isolate them from the males both socially and acoustically. The cage mates stayed together in the test cage during the whole testing period. However, just before and during a test, birds other than the test subject were moved to another soundproof chamber. A subject was isolated in the test cage 30 - 60 minutes before the start of song playback until the end of the test. All cage mates were returned to the test cage as soon as the test was finished.

In a single test, 5 songs (each consisting of 6 renditions) were played back in a random order (Fig. 1(b)). The inter-stimulus-interval was a random value ranging from 40 to 45 seconds. Thus, the duration of one test was approximately 450 seconds. Since some birds did not perform any CSDs in a test, the following criterion was set. We defined a valid test as one in which the subject expressed at least one CSD (in other words, a test was regarded invalid if she did not express any CSDs). We continued testing each female until 5 valid tests were obtained for that individual. One out of the 10 birds failed to meet the criterion due to health issues. We ceased testing that bird after 4 valid tests. The total number of tests required to reach the criterion varied between individuals (mean = 6.3 tests range = 5-9 tests). Each bird was tested once every 2 days to avoid habituation to stimuli. Thus, the test period for a subject lasted for 10-18 days, depending on the number of tests required. All the tests were conducted in the morning (7:00AM – 12:00PM).

### Recording and quantification of behavior

A web camera (BSW200MBK, Buffalo) recorded the movement and posture of subjects during the tests. The frame rate and sampling rate for video and audio recordings were 16 Hz and 44.1 kHz, respectively. The experimenter analysed the videos with the sound muted and had no access to any information regarding stimulus presentation order until the end of analysis, in order to minimize the effect of observer bias in behavioral quantification. For each stimulus in each test, we recorded the occurrence of CSDs performed during the song presentation period. A typical CSD could be recognized relatively easily: the bird usually titled her head and chest down, with her tail feather raised and quivering [24], as already described in other literature [22,25]. Across individuals, there were variations in the number of CSD bouts performed, the duration of bouts, and the posture displayed during CSDs. However, it was technically difficult to quantitatively code the response based on these features. Thus, we analyzed their response based on whether or not a bird performed CSDs in a trial (a song presentation in a test). For the analysis, data from all tests (both valid and invalid) in all subjects was used. The response rate for each stimulus was calculated as the number of tests in which a bird showed CSDs to that song divided by the total number of tests. We used this rate for description and figure illustration purposes but used the actual number of tests showing CSDs in the GLMM analysis detailed below.

### Evaluation of song similarity

If the development of female song preference depends on auditory experience with the father’s song, it can be expected that a bird’s response to a given song is predicted by the similarity between that song and the father’s song. To test this possibility, for each unfamiliar song stimulus, we calculated the acoustical and temporal similarity to the father’s song and analysed if these similarity indices predicted CSD frequency. For the evaluation of acoustical similarity, we applied the similarity measurement function (with asymmetric and time-courses mode) in Sound Analysis Pro 2011 [26] to the wave file data of song stimuli used in our song preference tests. For the similarity calculation, each song rendition was divided into 3 segments (duration of a segment = 2.00 ± 0.11 (mean ± *s*.*d*). seconds). Thus, the total number of files per stimulus was 18 (3 segments × 6 renditions). For every combination of segments between an unfamiliar song and the father’s song (18 × 18 = 324 combinations), the similarity value was computed and the mean value across these combinations was used as a representative value. To evaluate temporal similarity, we measured the song tempo, which is defined as the number of syllables divided by the duration from the first syllable onset to the last syllable offset. The mean song tempo of 6 renditions was used for each song. We then calculated the tempo difference (absolute value) between a given unfamiliar song and the father’s song.

### Statistical analyses

To statistically test if females showed more CSDs to the father’s song compared to unfamiliar songs, we fitted a generalized linear mixed model (GLMM) to a dataset [27] of all tests from all subjects. In the model, the dependent variable was the occurrence of CSDs in a given trial. The bird’s response was binarily coded as 0/1 (not performed/performed), respectively. The type of song stimulus (unfamiliar/father’s coded as 0/1) and song presentation order within a test (1-5) were the fixed effects of independent variables. Subject ID (10 levels), stimulus ID (19 levels), and test numbers (9 levels) were also included as random effects. The binomial distribution with a logit link function was specified to represent the probability distribution.

We next analyzed the possible effect of song similarity in the prediction of female response using a GLMM. In this analysis, we used trials of unfamiliar song playbacks from 7 birds that responded to at least 1 unfamiliar song [27]. The total number of trials was 176. The response rate to unfamiliar songs was low, and birds only responded in 17 trials. We fitted a model to test if acoustical and/or temporal similarity to the father’s song predicts female response to unfamiliar songs. The dependent variable was the occurrence of CSDs coded as 0/1 (not performed/performed), and binomial distribution with logit link function was specified. The SAP % similarity value, absolute value of the tempo difference, and song presentation order were the fixed effects of independent variables. Subject ID (7 levels), stimulus ID (17 levels), and test numbers (8 levels) were included as random effects.

GLMM estimation and statistical tests were conducted using R version 4.0.2 [28]. We used lme4 [29] and lmerTest [30] packages for GLMM analysis. In all the GLMM analysis, models were fitted using maximum likelihood estimation (Laplace approximation). For estimating the coefficients and their standard errors, *z*-value (Wald statistics) and *p*-value were calculated and reported in the results. For plotting data and fitting logistic regression curves, a Python-based package (SciPy version 0.19.0) was used.

## Results

We first examined if female Bengalese finches performed more CSDs in response to playback of the father’s song compared to unfamiliar conspecific songs. In all of the subjects we tested, the father’s song elicited more frequent responses compared to the 4 other songs. The mean response rate to the father’s song and the second-preferred unfamiliar song was 0.790 ± 0.209 and 0.223 ± 0.193 (mean ± *s*.*d*.), respectively (Fig. 2). The results of GLMM estimation indicated that the type of song stimulus (unfamiliar/father’s) was the strongest predictor of the birds’ response (*β ± s*.*e*. = 4.499 ± 0.544, *z* = 8.278, *p* < 0.001; Table 1). The presentation order within a test also significantly affected the birds’ response, although the estimated effect was relatively smaller than that of the song type (*β ± s*.*e*. = -0.346 ± 0.151, *z* = -2.297, *p* = 0.022).

**Table 1.** Effect of song type on females’ response.

| Independent variables | Estimates |  | *z*-value | *p*-value |
| --- | --- | --- | --- | --- |
| Fixed effects | coefficient | *s.e.* |  |  |
| Intercept | -1.957 | 0.598 |  |  |
| **song type (unfam/father = 0/1)** | **4.499** | **0.544** | **8.278** | **< 0.001** |
| **presentation order (1-5)** | **-0.346** | **0.151** | **-2.297** | **0.022** |
| Random effects | variance | *s.d.* |  |  |
| subject ID (10 levels) | 1.002 | 1.001 |  |  |
| stimulus ID (19 levels) | 0.000 | 0.000 |  |  |
| test number (9 levels) | 0.234 | 0.484 |  |  |

Estimated parameters of the GLMM are shown. Independent variables with a *p*-value less than 0.05 are indicated in boldface.

**Fig 2.**
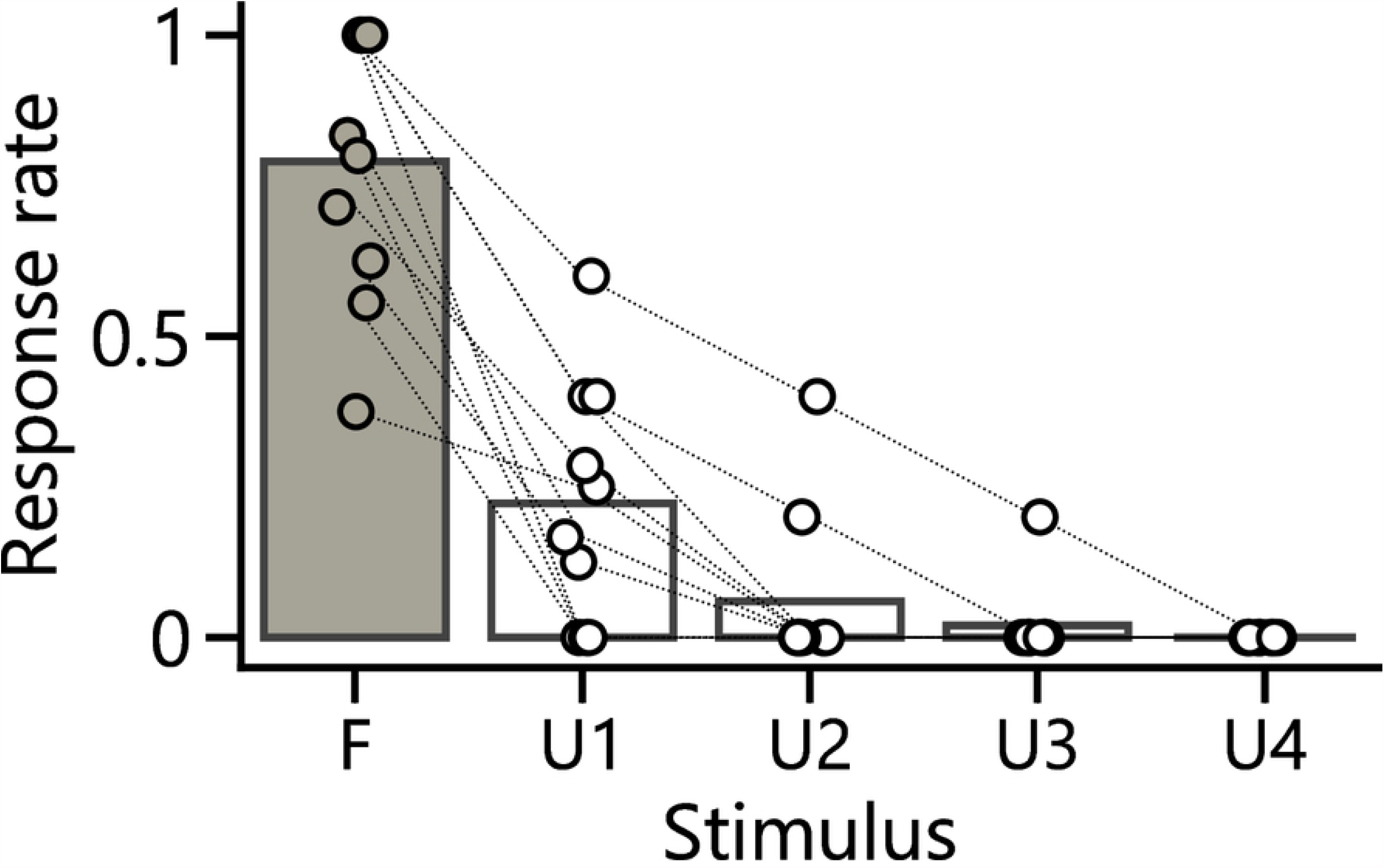
Females performed more CSDs to their father’s song. Response rate of CSDs to each stimulus (*n* = 10). Five songs are aligned in descending order of response rate. Dots connected with broken lines show individual data, while bars indicate the population mean.

We next conducted a post hoc analysis on whether similarity of a song stimulus to the father’s song predicted responses in the 7 birds that expressed CSDs to both song types (Fig. 3). Among the independent variables included in the GLMM, presentation order within a test was the strongest predictor of the response (Table 2). As with the result in Table 1, a song was more likely to elicit CSDs when it was presented earlier in a test (Fig. 3 (c)). Although there was a tendency for unfamiliar songs to elicit more frequent CSDs when these songs had greater similarity to the father’s song (higher SAP % similarity, lower tempo difference, Fig. 3(a), (b)), this result was not statistically significant (Table 2). The overall results imply that female Bengalese finches in our study did recognize their father’s song as an attractive courtship signal, but we cannot not conclude whether their mate choice is affected by song features that are shared with their father’s song.

**Table 2.**
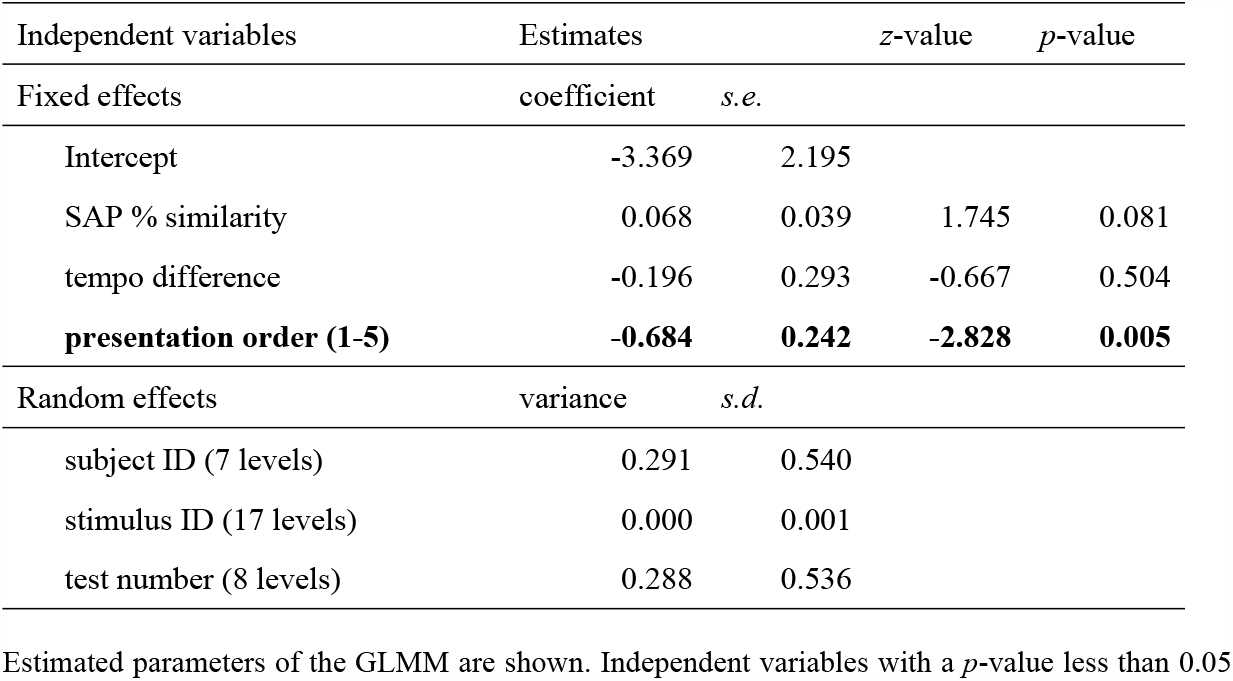
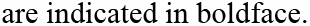
Effect of song similarity on females’ response to unfamiliar songs.

| Independent variables | Estimates |  | *z*-value | *p*-value |
| --- | --- | --- | --- | --- |
| Fixed effects | coefficient | *s.e.* |  |  |
| Intercept | -3.369 | 2.195 |  |  |
| SAP % similarity | 0.068 | 0.039 | 1.745 | 0.081 |
| tempo difference | -0.196 | 0.293 | -0.667 | 0.504 |
| **presentation order (1-5)** | **-0.684** | **0.242** | **-2.828** | **0.005** |
| Random effects | variance | *s.d.* |  |  |
| subject ID (7 levels) | 0.291 | 0.540 |  |  |
| stimulus ID (17 levels) | 0.000 | 0.001 |  |  |
| test number (8 levels) | 0.288 | 0.536 |  |  |

Estimated parameters of the GLMM are shown. Independent variables with a *p*-value less than 0.05
are indicated in boldface.

**Fig 3.**
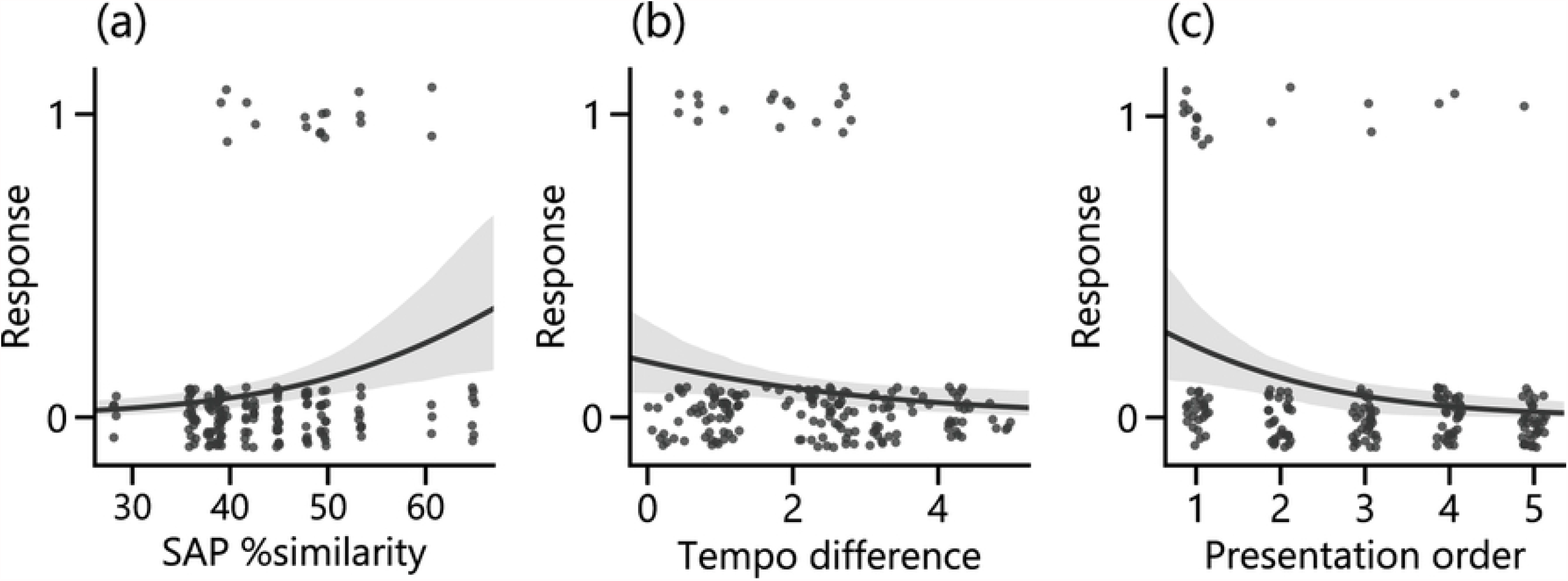
Analysis of song similarity. CSD response was plotted against the **(a)** acoustical similarity (SAP % similarity), **(b)** temporal similarity (absolute value of tempo difference) of a given unfamiliar song to the father’s song, or **(c)** presentation order within a test (1-5). In all the panels, 0 and 1 on the vertical axis means ‘not performed’ or ‘performed’, respectively. Each data point corresponds to one trial. Thus, there are 176 data points from 7 birds in total (17 points are plotted on ‘performed’). The lines and surrounding bands are the logistic regression curve with 95% confidential interval computed by bootstrapping method (iteration *n* = 1000).

## Discussion

Previous studies in Bengalese finches and zebra finches have found that female birds develop a long-lasting preference for their father’s song [10–15]. From a functional viewpoint, however, it was not clear if such a preference is an expression of sexual motivation in females. In the current study, we tested whether female Bengalese finches are sexually attracted to their father’s song by measuring CSDs in response to song playbacks. We found that birds showed greater frequency of CSDs to the father’s song than to other unfamiliar conspecific songs (Fig. 2), which suggests that females are sexually motivated when exposed to the father’s song.

The song preference observed in our study might be regarded as an expression of sexual imprinting on the father’s song. This possibility has already been referred to in an early study of captive zebra finches, although there has not been a conclusive experimental demonstration of this to date [10]. Likewise, field research in Darwin’s finches (*Geospiza*) reported female mate preference consistent with sexual imprinting on the father’s song [31]. However, the effect of experience with the father’s song on later mate preference, if any, may depend on the species. In canaries (*Serinus canaria*), for example, adult females rather disfavor their foster father’s song, which likely helps the species avoid inbreeding [20]. On the other hand, it can be speculated that learning the father’s song as a sexually attractive signal may help other species avoid outbreeding or hybridization [32–35]. Whether females of a species favor or disfavor incestuous signals might depend on the dispersal patterns and breeding ecology of that species [16,36,37].

To examine if female Bengalese finches are sexually imprinted on their father’s song, however, more precise control of rearing conditions is necessary. For example, our experimental design could not exclude the possibility that song preference was somehow inherited from parents to daughters, as we used female finches that were cared for by their genetic parents. In addition, because the subjects were housed with parents for a relatively long time (until about 120 dph), it is also possible that such a long-time interaction with the father led to a particularly robust response selectivity to conspecific songs. To avoid these confounding factors and better understand the process of preference development, cross-fostering experiments and/or manipulation of the period of exposure to (foster) father’s song will be necessary in the future. Moreover, although we found that songs with higher similarity to the father’s song elicited more CSDs, this tendency was not statistically significant. In testing the effect of similarity on female response to unfamiliar songs, the small number of trials available for analysis was a limitation (Fig. 3). Manipulation of stimuli based on song similarity or a particular song feature, rather than a post-hoc analysis as in this study, may enable a more detailed analysis on the effect of auditory experience on song perception.

Regardless of the developmental mechanism of preference for the father’s song, our results together with previous findings [14,15] imply the importance of considering individual variations in female song preference in the Bengalese finch. In previous studies on this species, researchers have explored acoustical and temporal song characteristics that are preferred across individuals [14,38–40]. For example, in a call-back assay experiment where song tempo or pitch were manipulated, the authors found that the majority of females preferred faster songs while changes in pitch were not a good predictor of female response [38]. There are other studies that are especially relevant to the characteristics of Bengalese finch song and its evolutionary process. The Bengalese finch is a domesticated strain of its wild ancestor, the white-rumped munia (*Lonchura striata*), and the transition probability of Bengalese finch song elements is known to be more complex than that of munias [41,42]. Thus, it is hypothesized that sexual selection contributed to an increase in sequential complexity across generations. If this is the case, one would predict that that females should possess a preference for such complexity [42]. However, the results of studies have been mixed: although some females did show greater responses to more complex songs (measured by operant conditioning or nest building behavior), there were individual variations in the preference for or sensitivity to the sequential complexity of song [14,39,40]. Individual differences in song preference were also reported in another study [22], but it was not known what factor(s) might explain such variation. Our results here suggest that developmental song experience may account for this issue. Among different studies that used different behavioral indices, there is general agreement that as a population, female birds tended to prefer their father’s song [14,15]. We propose that consideration of the early-life social environment of females is important in re-interpreting previous findings and designing new experiments.

## Acknowledgements

We thank Dr. Maki Ikebuchi for assistance in bird breeding, and Dr. Chihiro Mori for instruction of hormone implantation. We are grateful to Dr. Beth A. Vernaleo for her careful proof reading of this paper.

